# Teacher-student neural coupling during teaching and learning

**DOI:** 10.1101/2020.05.07.082958

**Authors:** Mai Nguyen, Ashley Chang, Emily Micciche, Meir Meshulam, Samuel A. Nastase, Uri Hasson

**Affiliations:** Department of Psychology, Princeton University, Princeton, NJ 08540; Department of Psychology, Vanderbilt University, Nashville, TN 235; Princeton Neuroscience Institute, Princeton University, Princeton, NJ 08540

**Author notes:** **Corresponding authors:** Mai Nguyen,; Uri Hasson.

## Abstract

Human communication is remarkably versatile, enabling teachers to share highly abstracted and novel information with their students. What neural processes enable such transfer of information across brains during naturalistic teaching and learning? Here, we show that during lectures, wherein information transmission is unidirectional and flows from the teacher to the student, the student’s brain mirrors the teacher’s brain and that this neural coupling is correlated with learning outcomes. A teacher was scanned in fMRI giving an oral lecture with slides on a scientific topic followed by a review lecture. Students were then scanned watching either the intact lecture and review (*N* = 20) or a temporally scrambled version of the lecture (*N* = 20). Using intersubject correlation (ISC), we observed widespread teacher-student neural coupling spanning sensory cortex and language regions along the superior temporal sulcus as well as higher-level regions including posterior medial cortex (PMC), superior parietal lobule (SPL), and dorsolateral and dorsomedial prefrontal cortex. Teacher-student alignment in higher-level areas was not observed when learning was disrupted by temporally scrambling the lecture. Moreover, teacher-student coupling in PMC was significantly correlated with learning outcomes: the more closely the student’s brain mirrored the teacher’s brain, the more the student improved between behavioral pre-learning and post-learning assessments. Together, these results suggest that the alignment of neural responses between teacher and students may underlie effective communication of complex information across brains in classroom settings.

**Significance statement:** How is technical, non-narrative information communicated from one brain to another during teaching and learning? In this fMRI study, we show that the DMN activity of teachers and students are coupled during naturalistic teaching. This teacher-student neural coupling emerges only during intact learning and is correlated with learning outcomes. Together, these findings suggest that teacher-student neural alignment underlies effective communication during teaching.

## Introduction

Humans have a unique ability to share knowledge of the world with each other via communication. Often times, communication takes the form of sharing narratives or recalling memories in social contexts where interlocutors have a common background and shared knowledge. However, in other contexts, such as in teaching and learning, human communication is often asymmetric and involves the transfer of novel, non-social information from an expert (teacher) to a novice (student). What neural processes enable transfer of information across brains during naturalistic teaching and learning of complex, abstract information?

Verbal communication of social narratives elicits correlated neural responses between speakers and listeners in regions overlapping with the default mode network (DMN; Stephens et al., 2010; Dikker et al., 2014; Silbert et al., 2014; Zadbood et al., 2017). This “speaker-listener coupling” is thought to be driven by shared understanding of the narrative: speaker-listener coupling in DMN is observed only during comprehensible communication and is correlated with listener comprehension (Stephens et al., 2010; Silbert et al., 2014). Moreover, shared DMN responses among listeners is sensitive to background context or knowledge that enables similar comprehension of interpretation of the narrative. Subjects who receive contextualizing background information show more strongly correlated neural responses in DMN in response to ambiguous narratives than subjects lacking this information (van Kesteren et al., 2010; Ames et al., 2014; Chen et al., 2015; Oren et al., 2017). Similarly, subjects who are biased by context to interpret the stimulus in same way have more similar DMN responses with each other than with subjects who are biased towards a different interpretation (Yeshurun et al., 2017; Chien and Honey, 2020).

However, during communication of non-narrative, technical information, individuals may lack a common context or background for communicating effectively. Teaching is therefore a means for establishing a shared common ground for understanding new information. The teacher has abstract, complex information represented in their brain, which must be transmitted to students, who may initially lack the context or schema required to understand this novel information. Here, we suggest that the process of establishing this shared knowledge via teaching is reflected in coupled neural processes between the teacher and students in regions of the DMN. While previous fMRI studies have identified speaker-listener alignment in the communication of narratives, these findings have not been extended to the communication of non-narrative, technical information. Teacher-student neural coupling during non-narrative communication has been measured using EEG and fNIRS (Zheng et al., 2018; Pan et al., 2020; Bevilacqua et al., 2019, but our use of fMRI imaging in the present work allow for full-brain coverage with high spatial resolution. EEG in particular does not allow for measurements from regions of interest in the DMN, including posterior medial cortex and medial prefrontal cortex.

Here, we extend prior research on the neural basis of real-life learning by using fMRI to investigate teacher-student alignment during teaching and learning of complex scientific material. We scanned a teacher giving an extended, 32-min lecture on a scientific topic followed by a 6-min review of the material. Students were then scanned while watching the video lessons and taking pre- and post-lesson quizzes to assess learning. Based on previous work, we predicted watching engaging video lessons would elicit shared, or correlated, neural responses among students in regions ranging from early sensory cortex to high-level regions of the default mode network (DMN). We additionally predicted that the shared neural response among students in high-level regions is driven by activity in the teacher’s brain, and that this coupling of the teacher and students’ brains will correlate with learning outcomes, particularly doing the review session. Finally, because the main lesson serves to create a shared knowledge between teachers and students, we predicted that teacher-student coupling will be greater during the review than the lesson.

## Methods

### Subjects

One teacher (author M.N.; age: 27; female) was scanned using fMRI while giving a verbal lecture and review with accompanying slides. Forty-eight subjects with normal hearing and normal or corrected-to-normal vision participated in the experiment as students. The data from four subjects were excluded from analysis for excessive motion during scanning (>3mm), two for falling asleep during scanning, and two for significant pre-existing knowledge of the lesson material (scored >=80% on the pre-lesson test). This left 20 subjects (ages 19–34, mean = 22.2 years; 13 female) who participated in the Intact Learning condition and 20 subjects (ages 18-26, mean=19.65 years; 10 female) who participated in the Scrambled control condition. All experimental procedures were approved by the Princeton University Internal Review Board, and all subjects provided written, informed consent.

### Experimental design

#### Stimuli

The teacher developed a 32-min Lesson and a 6-min Review of the lesson, both with slides (Figure 1A). In order to engage the subjects with the lesson, we selected fMRI as the lesson topic, covering an introduction to fMRI, fMRI physics, action potentials, and the biological basis for the blood-oxygen-level-dependent (BOLD) signal (Figure 1B). The teacher was then scanned using fMRI while giving the Lesson and Review. The teacher repeated the same Lesson and Review five times across three different scanning sessions (Lesson duration: mean = 32:20 min, range = 31:29–34:29 min; Review duration: mean = 5:58 min, range = 5:54–6:13 min). During teaching, slides were presented using PsychoPy 2 (Peirce et al., 2019) and presented via an LCD projector on a rear-projection screen mounted in the back of the scanner bore. The teacher was able to view slides through a mirror mounted on the head coil and advance slides using a button box. The teacher’s speech was recorded using an MRI-compatible microphone with online sound canceling (FOMRI III; Optoacoustics Ltd).

**Figure 1.**
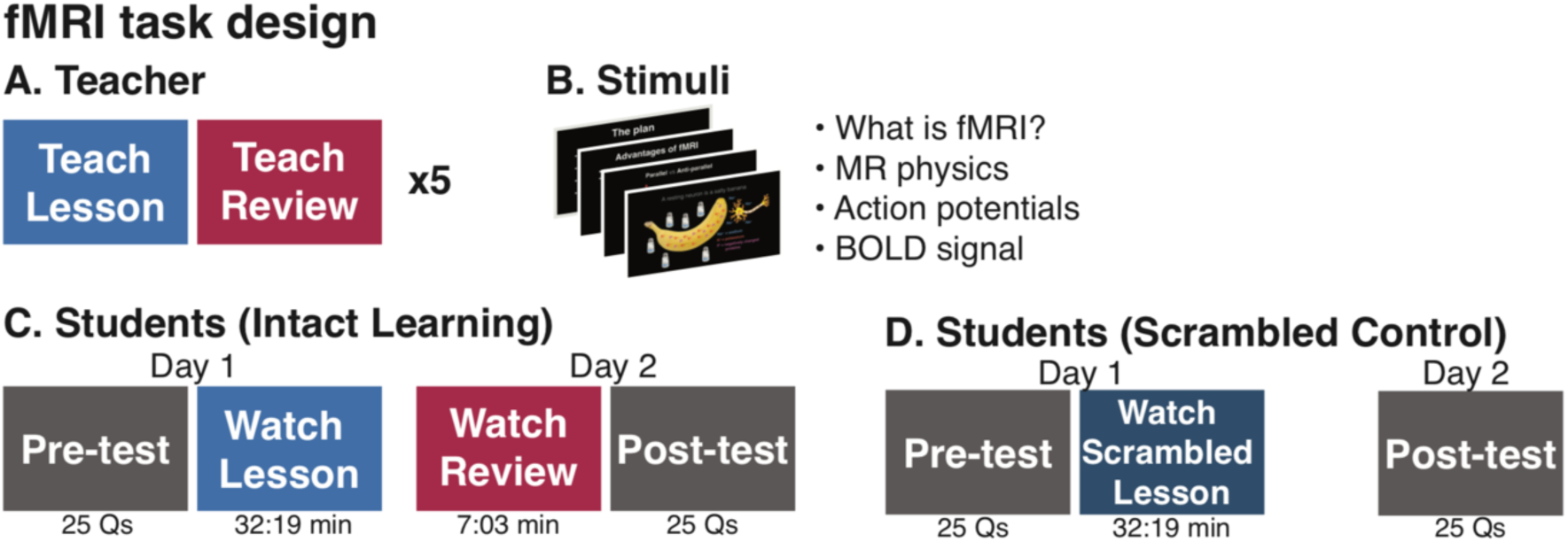
fMRI task design. (A) A teacher was scanned in fMRI giving a 32-min lessonand a 7-min review with slides, repeating each five times in the scanner while her voice were recorded. (B) The lecture and review introduced fMRI concepts. (C) Students in the Intact Learning condition took a Pre-test quiz and watched the Lesson in the scanner. On the following day, they were scanned watching the Review and then completed a Post-test quiz. (D) Subjects in the Scrambled condition took the same Pre- and Post-test quiz, but watched a scrambled version of the Lesson.

Following the teacher scans, the audio recordings were further denoised using Adobe Audition, transcribed and timestamped, and then aligned with slide presentation. The highest-quality recordings were then selected to present to students. The Lesson presented to students was 32:19 min, and the Review was 6:13 min (see SI for videos). To minimize transient, non-selective responses that occur at the abrupt onset of a stimuli, the lectures were preceded by the same unrelated 37 s movie clip followed by a 6 second fixation cross. Videos ended with another 6 seconds of fixation cross. The unrelated movie clip and periods of fixation were cropped from subsequent analyses.

Finally, the teacher developed a 25-question, multiple choice quiz to assess learning. The same quiz was administered before and after learning (Pre-test, Post-test).

#### Student sessions: Intact Learning

Student subjects completed the main experiment on two consecutive days. In the Intact Learning condition, on Day 1, subjects (*N* = 20) took the multiple choice quiz (Pre-test) assessing their pre-existing knowledge of the lesson topic. They were then scanned in fMRI watching the 32-min Intact Lesson. On Day 2, subjects watched the 6-min Review, completed additional scans that were not analyzed here, and then answered the same 25-question quiz (Post-test). Stimuli were presented and viewed in an identical setup as during the teacher sessions. Audio was played through MRI-compatible insert earphones (Sensimetrics models S14 and S15). At the start of each scanning session, a short unrelated audio clip of the teacher was played during a scan to calibrate audio volume. Following scanning, students were asked to rate how closely they attended to the video on a scale of 1–5 (1 = not at all attentive, 5 = extremely attentive).

#### Student sessions: Scrambled Lesson control

To test that neural coupling between the teacher and students reflected shared processing of lesson content, rather than shared perception of low-level stimulus features, a second sample of subjects (*N* = 20) was scanned while watching a Scrambled Lesson control. In this control, the 32-min Intact Lesson was scrambled at the sentence level, while the text on the slides was scrambled at the word level. Subjects in the Scrambled Lesson control also completed the experiment on two consecutive days: on Day 1, students took the Pretest and were scanned watching the Scrambled Lesson and then made the same ratings as Intact Learning subjects; on Day 2, students took the Posttest outside of the scanner.

### MRI acquisition

Subjects were scanned in a 3T Siemens Magnetom Skyra scanner at the Princeton Neuroscience Institute using a 64-channel head/neck coil (Siemens). During functional scans, volumes were acquired using a T2*-weighted gradient-echo echo-planar pulse sequence (TR 2000ms; TE 28ms; voxel size 3×3×3mm with 38 slices; flip angle 80°; FOV 192×192 mm^2^; matrix size 64×64; slice orientation axial; anterior-to-posterior phase encoding; interleaved slice acquisition; iPAT GRAPPA 2) with whole-brain coverage. For both Intact and Scrambled Lesson, total scan duration was 966 volumes (32:12min). For the Review, total scan duration was 207 volumes (6:54min). A high-resolution anatomical image was collected using a T1-weighted MPRAGE pulse sequence (TR 2300ms; TE 2.98ms; TI 900ms; voxel size 1×1×1mm with 176 slices; flip angle 9°; FOV 256×256 mm^2^; slice orientation axial; no fat suppression; total scan duration 5:20 min).

### MRI data analysis

#### Preprocessing

MRI data were preprocessed using FSL 5.0 and FEAT 6.0 (FMRIB, Oxford; (Jenkinson et al., 2012) including motion correction, linear trend removal, high-pass filtering (cutoff: 140s or ~.007 Hz), and spatial smoothing with a 6-mm FWHM Gaussian kernel. Motion correction was performed using FSL’s MCFLIRT with 6 degrees of freedom. Subjects with excessive head motion (>3 mm absolute displacement) were discarded from the analysis. Functional data were registered to high-resolution structural images and then to 3-mm MNI standard space using FSL’s FLIRT with 12 degrees of freedom. Preprocessed data were z-scored over time. All analyses were conducted in volume space using custom MATLAB and Python scrips and visualized using nilearn (https://nilearn.github.io; (Abraham et al., 2014) and NeuroElf (https://neuroelf.net).

#### Temporal interpolation procedure for Teacher scans

Scans of the teacher giving the Lesson and Review were separately averaged to provide a more reliable measure of the Teacher’s neural responses. The Teacher was extensively trained to reproduce the same Lesson and Review with high fidelity in timing and word use. To account for natural and minor variation in timing, the recorded Lessons and Reviews were timestamped at the approximate sentence level (137 segments, mean length = 13.78 sec, SD = 4.11 sec). The teacher’s neural response was then linearly interpolated within each timestamped sentence in order to align stimulus content over time relative to the actual recordings presented to the student subjects. The interpolated teacher neural responses were then averaged and used in subsequent analyses.

#### Unscrambling procedure for Scrambled Lesson control

The order of sentences in the Scrambled Lesson were randomized, so in order to compute coupled neural responses between the Teacher and Student Scrambled Lesson, the Scrambled Lesson response timeseries were reordered to match the temporal order of the Intact Lesson. Following Lerner et al (Abraham et al., 2014), sentence-level timestamps were shifted by 6 seconds (3 TRs) to account for the hemodynamic response function, and then the response timeseries were segmented by sentence and reordered. Sentences less than six seconds long were removed from analysis. The resulting reordered timeseries was used in the Teacher-Student “Unscrambled” Lesson analyses described below. The full, original scrambled timeseries was used in Student-Student Scrambled analyses.

#### Intersubject correlation (ISC)

Shared neural responses among subjects were measured using intersubject correlation (ISC; Hasson et al., 2004; Nastase et al., 2019). Four ISC analyses were conducted: (1) among students in the Intact Lesson and Review (Student-Student), (2) among the scrambled students in the Scrambled Lesson control (Student-Student Scrambled), (3) between the teacher and the students in the Intact Lesson and Review (Teacher-Student), and (4) between the teacher and the students in the Scrambled Lesson control (Teacher-Student Unscrambled). ISC was calculated at each voxel within an anatomically-defined gray matter mask. For within-group analyses among students (comparisons 1 and 2), ISC was calculated using the leave-one-out approach: each student’s response time course was correlated to the average response time course of the remaining (*N* – 1) students at each voxel. ISC was separately calculated among the students in the Intact Learning condition for both Lesson and Review and among the students in the Scrambled Control condition for the Scrambled Lesson only. The same analysis was repeated for the between-group Student and Teacher analyses (comparisons 3 and 4). In these analyses, ISC was calculated by correlating each student’s response time series to the average Teacher response time series.

Statistical significance of ISC was assessed using a bootstrap hypothesis test following Chen et al. (2016) and Nastase et al. (2019). In each iteration of the bootstrap, we randomly sampled *N* = 20 subject ISC values with replacement then computed the mean of the bootstrap sample. We repeated this procedure 10,000 times, producing a bootstrapped distribution around the mean ISC value across subjects. We then subtracted the observed mean ISC value from the bootstrap distribution to create a null distribution, and used this null distribution to calculate *p-*values. We corrected for multiple comparisons across voxels by controlling the false discovery rate (FDR) (Benjamini and Hochberg, 1995) at *q* = .05.

#### ISC in Lesson versus Review

We predicted that Teacher-Student coupling during the review will be greater than during the lesson as the initial lesson serves to build a shared knowledge background. To test this prediction, we directly compared Teacher-Student and Student-Student ISCs in the Intact Lesson and Review conditions using paired *t*-tests (two-tailed) in every region of a 61 region parcellation comprising regions of the brain that respond reliably to naturalistic audiovisual stimuli (Regev et al., 2018). Statistical significance was assessed using a sign test: in each permutation, the sign of the difference in the two conditions was randomly flipped, and the *t-*value was calculated under the null hypothesis of no systematic difference between conditions. This procedure was repeated 10,000 times to produce a permutation-based null distribution, which was used to calculate *p*-values. We corrected for multiple comparisons using FDR (*q* =.05).

#### Correlation with learning

To test the prediction that the level of shared neural responses will be correlated with learning, we first calculated a normalized measure of learning outcome: normalized score = (Posttest – Pretest) * mean(Posttest + Pretest). This measure incorporates both improvement in score and total score, balancing the amount a student learns with an absolute measure of how much they know. Two students may improve by the same amount, but a student who has a higher total score, indicating greater understanding of the material, will have a higher normalized score. We correlated both Student-Student ISC and Teacher-Student ISC in the Intact Lesson and Review in each of the 61 ROIs with the normalized score. Statistical significance of the ISC-normalized score correlation was assessed using a permutation test. In each permutation, the ISC values were randomly shuffled across subjects and correlated with the normalized scores. This procedure was repeated 10,000 times, producing a permutation-based null distribution (under the null hypothesis of no systematic relationship between ISC values and learning scores across subjects). We again corrected for multiple comparisons using FDR (*q* = .05).

#### Lagged correlation

Previous research has suggested that during verbal communication, the listener’s neural response follows the speaker’s neural response with a several second lag in certain brain areas. We therefore tested for a lag between student and teacher neural responses during the Intact Lesson in two parcels, posterior cingulate cortex (PCC) and V1+, by shifting the Students’ timeseries by −10 to +10 TRs relative to the teacher’s timeseries and calculating ISC at each lag. The same analysis was repeated for Student-Student ISC in the Intact Lesson by shifting each subject relative to the average of others.

## Results

### Behavioral results

Subjects in the Intact Learning condition improved significantly from the Pre-test to the Post-test Quiz (*t*(19) = 10.35, p < .001), scoring on average 48.8% (SD = 12.2%) on the Pre-test and 87.2% (SD = 17.5%) on the Post-test (Figure 2A). All subjects in the Intact Learning condition improved their score. Subjects in the Scrambled Lesson control condition also significant improved their scores (Pre-test: mean = 47.4%, SD = 1 3.3%; Post-test: 59.4%, SD = 14.1%; *t*(19) = 3.47, *p* = .003), but by significantly less than the subjects in the Intact Learning condition (*t*(38) = 5.21, *p* < .001). The two groups did not differ significantly in Pre-test scores (*t*(38) = .345, *p* = .73), indicating a similar baseline of prior knowledge.

**Figure 2.**
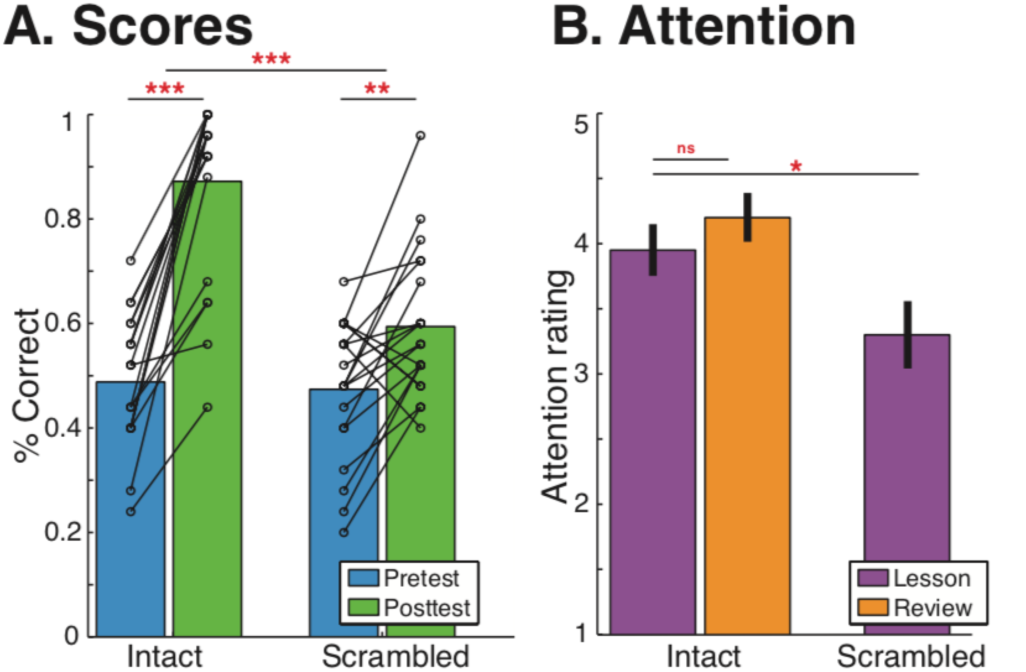
Behavior. (A) Students in both the Intact and Scrambled conditions significantly improved their scores from Pre-test to Post-test (Intact: t(19)=10.35, p<.00001; Scrambled: t(19)=3.47, p=.0026), but Students learned significantly more in the Intact condition (t(38)=5.21, p<.00001). (B) On average, students were very attentive during the videos, but students in the Intact Lesson were more attentive than students in the Scrambled lesson (t(38)=2.27, p=.029). ***p<.00001, **p<.01, *p<.05

Subjects in both conditions were attentive throughout the video lessons (Figure 2B). On a five-point scale (1 = not at all attentive, 5 = extremely attentive), subjects in the Intact Learning condition rated their attention during the Lesson on average to be 3.95 (SD = .76) and during the Review to be 4.20 (SD = .70). There was no significant difference between attention during the Lesson and the Review (*t*(19) = 1.56, *p* = .14). During the Scrambled Lesson control, subjects rated their attention as 3.30 (SD = 1.03), which is significantly lower than the subjects in the Intact Lesson (*t*(38) = 2.27, *p* = .029).

### fMRI results

#### Aligned neural responses among students during learning

We first identified regions of the brain that are significantly correlated responses across students during the Lesson and Review in the Intact Learning condition. Consistent with previous work using audiovisual narrative stimuli, we observe widespread significant ISC in during both the Lesson and Review (*q* < .05, FDR corrected voxelwise; Figure 3A) throughout much of visual cortex, auditory and linguistic regions from early auditory cortex (A1+) to superior temporal gyrus (STG) and middle temporal gyrus (MTG), and extending into higher-order DMN regions including posterior medial cortex (PMC), bilateral dorsolateral prefrontal cortex (DLPFC), and right medial prefrontal cortex (mPFC). However, unlike previous work on narrative processing (e.g. Hasson et al., 2008; Lerner et al., 2011; Yeshurun et al., 2017; Nguyen et al., 2019), we also observe strong ISC in bilateral superior parietal lobule (SPL). There were no significant differences between Student-Student ISC during the Lesson versus Review.

**Figure 3.**
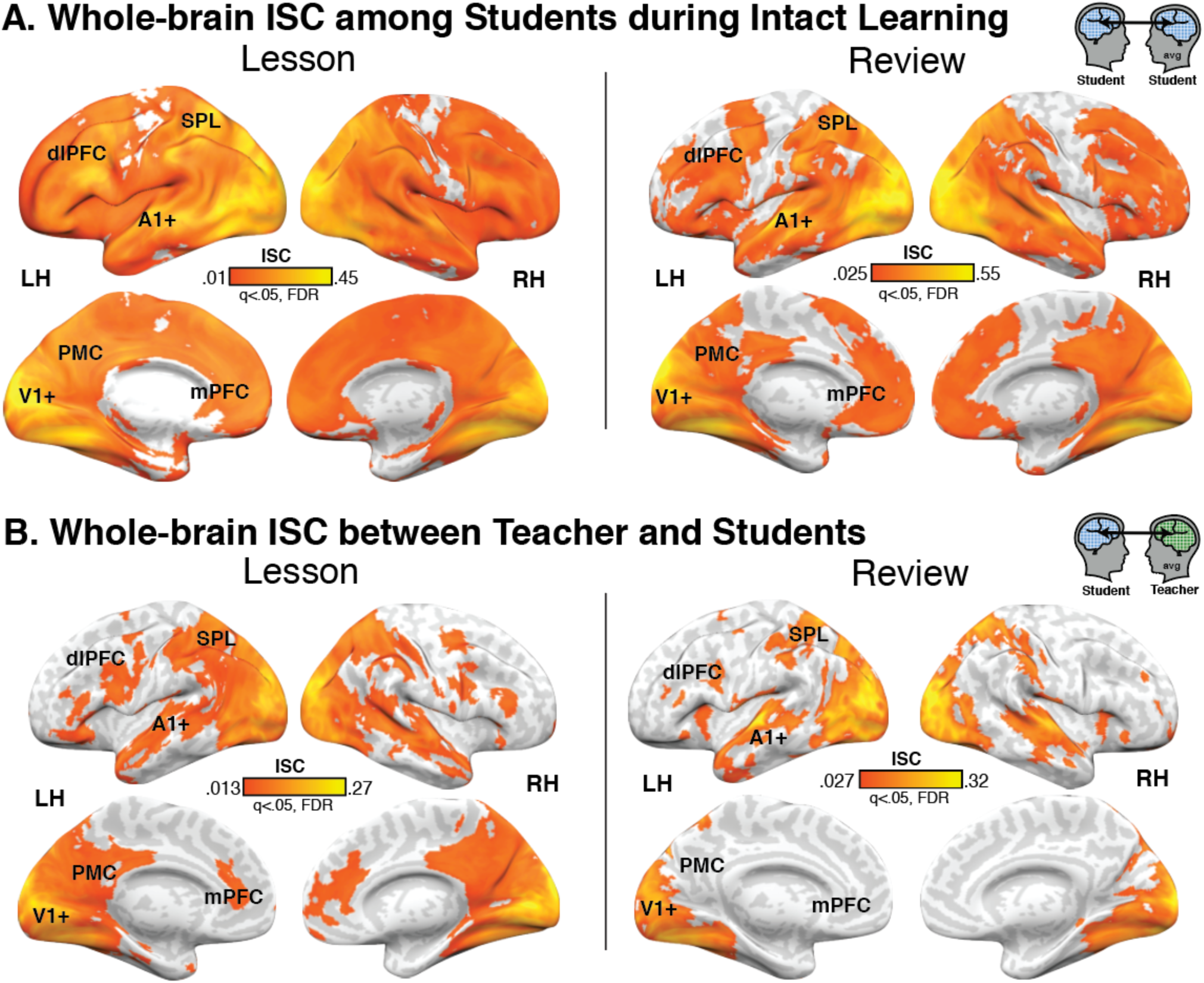
ISC. (A) Students in the Intact Learning condition have significant ISC throughout the cortex in both the Lesson and Review, with the strongest ISC in visual and auditory cortex, superior and middle temporal gyrus, bilateral SPL, bilteral DLPFC, and right mPFC. (B) Student neural responses are coupled to the teacher’s response in similar regions. dlPFC = dorsolateral prefrontal cortex, SPL = superior parietal lobule, PMC = posterior medial cortex, mPFC = medial prefrontal cortex (non-parametric bootstrap hypothesis test; q<.05, FDR corrected).

#### Teacher-student neural coupling during teaching and learning

We next calculated ISC between the Teacher and Student neural responses in the Intact Learning condition to identify regions of the brain that are coupled during teaching and learning (Figure 3B). Similarly to speaker-listener coupling during narrative storytelling, significant Teacher-Student ISC was observed in early sensory cortices, linguistic and extralinguistic regions including STG, MTG, and temporal pole, and DMN regions including PMC, dlPFC, and mPFC. Unlike coupling during narrative storytelling, we additionally observe significant ISC in bilateral SPL and notably weak ISC in bilateral TPJ. Again, there were no significant differences between Teacher-Student ISC during the Intact Lesson versus Review.

The lagged correlation analysis does not indicate a lag between the Teacher and Students’ neural responses. In both V1 and PCC, Student-Student and Teacher-Student ISC peaks at lag = 0, suggesting that fluctuations in neural response in the Students and Teacher was locked to the slides progression as the lectures unfold over time.

#### Teacher-student ISC emerges in DMN only during intact learning

Shared neural responses between the Teacher and Students could be driven by shared sensory input rather than shared high-level understanding of the lesson content as the Teacher and Students view and hear the same visual and auditory stimuli. To dissociate content-related shared responses from shared sensory responses, we scanned an additional group of students who watched a temporally scrambled version of the Lesson. This manipulation preserves the low-level visual and auditory properties of the stimulus while disrupting conceptual understanding and reducing learning (Figure 2A). The neural response of students in the Scrambled Lesson control were “unscrambled” to reorder events in the same order as the original Intact Lesson and then correlated with the Teacher’s neural response.

Consistent with the hypothesis that Teacher-Student coupling in high-order areas reflects shared understanding, not just shared stimulus processing, we observed significant Teacher-Student ISC in DMN regions only in the Intact Learning conditions (Figure 4). These regions include PMC, bilateral angular gyrus, bilateral temporal poles, mPFC, and lateral PFC. Teacher-Student Unscrambled ISC largely overlapped with the ISC in the Intact Lesson in visual areas, extending from V1+ to ventral-temporal cortex and dorsal occipital regions. In addition, however, there was significant Teacher-Student Unscrambled ISC in attention-related regions of dorsal PFC that did not overlap with the Intact Lesson ISC map.

**Figure 4.**
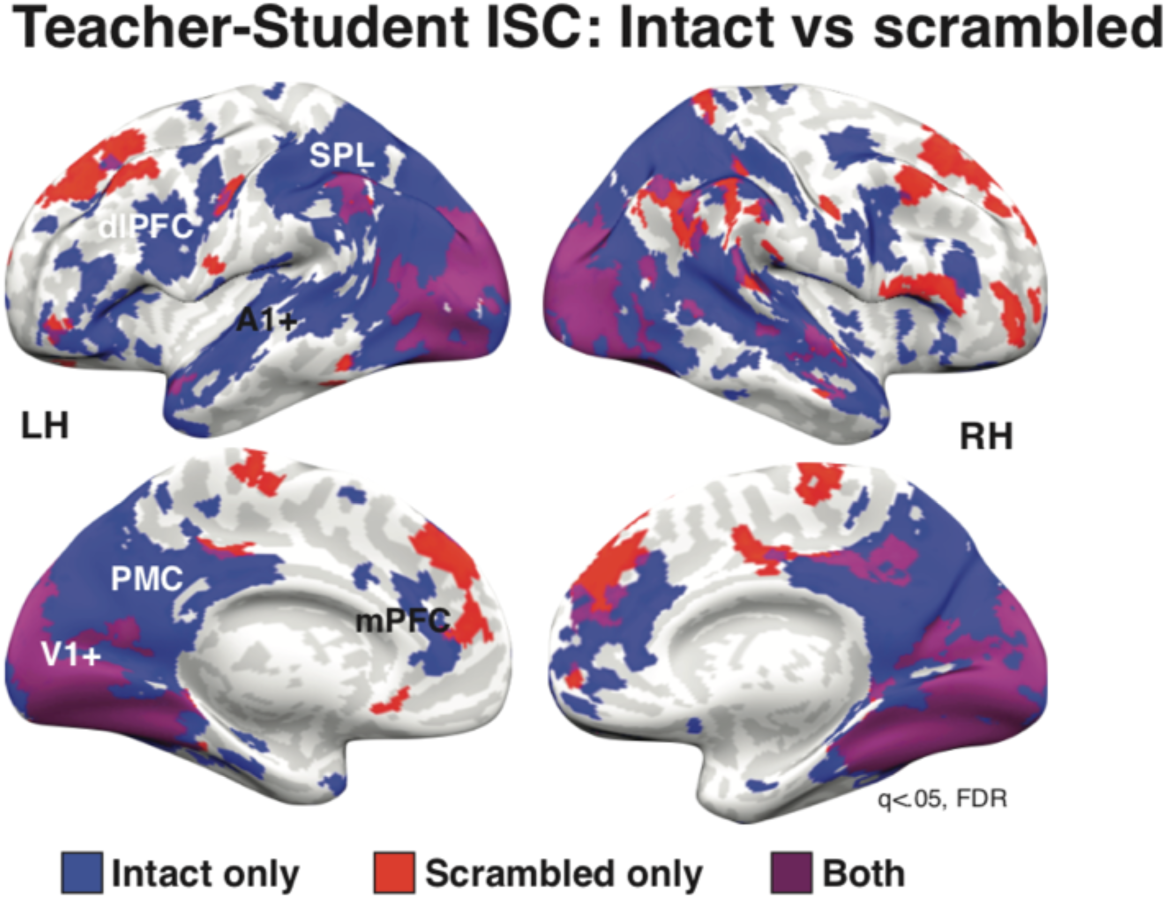
Scrambled vs intact ISC. ISC in both the Teacher-Intact Student Intact and Teacher-Scrambled Student conditions were significant in visual cortex. However, only the Intact condition showed significant ISC in high-level DMN regions, including PMC, MPFC, and DLPFC. Surprisingly, the Scrambled condition also showed significant ISC in dorsal PFC. dlPFC = dorsolateral prefrontal cortex, SPL = superior parietal lobule, PMC = posterior medial cortex, mPFC = medial prefrontal cortex (q<.05, FDR corrected).

#### Teacher-Student ISC correlates with learning

If Teacher-Student coupling reflects shared understanding of novel information, we also expect that coupling in high-level regions, but not low-level sensory regions, will be related to learning outcomes. We therefore correlated both Student-Student and Teacher-Student ISC with a measure of learning outcomes (improvement in score weighted by total score) across each of 61 areas from an independent parcellation (Regev et al., 2018). The analysis revealed seven regions with a significant correlation between Teacher-Student ISC during the lesson and learning outcomes (Figure 5, FDR corrected). Five regions were located primarily in posterior medial cortex: posterior cingulate cortex (PCC; *r* = .663, *p* = .002), right precuneus (rPCUN; *r* = .626, *p* = .003), left precuneus (lPCUN; *r* = .751, *p* < .001), superior occipital gyrus (SOG; *r* = .634, *p* = .003), and dorsal precuneus (dPCUN; *r* = .755, *p* < .001). Two regions are in high-order visual cortex: left lateral occipital complex (lLOC; *r* = .633, *p* = .003) and right hV4 (rhV4; *r* = .619, *p* = .004). Notably, these areas largely overlap with regions that only show significant Teacher-Student coupling in the Intact Learning condition.

**Figure 5.**
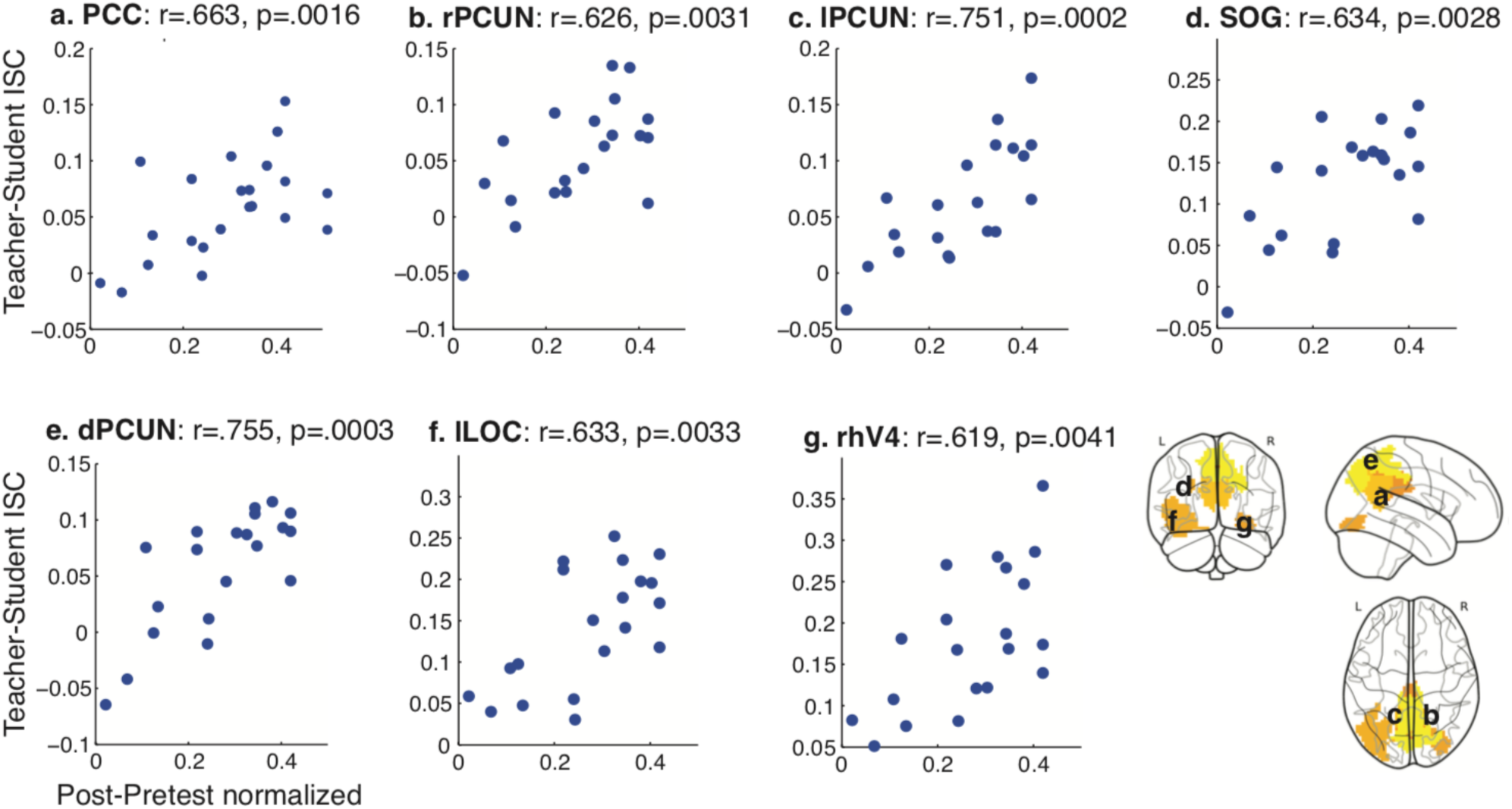
ISC correlation with behavior. Teacher-student ISC is correlated with normalized improvement in quiz score (Post-Pre/ (1/avg(Post+Pre)) in five ROIs, primarily in posterior medical cortex (q<.05, FDR corrected). PCC = posterior cingulate cortex, rPCUN = right precuneous, lPCUN = left precuneous, SOG = superior occipital gyurs, dPCUN = dorsal precuneous, lLOC = left lateral occipital complex, rhV4 = right human V4. ROIs from 61 ROI parcellation (Regev et al., 2018).

While the correlation between Student-Student ISC and improvement in quiz score showed a similar pattern of effects, no region passed FDR correction. There was no relationship between learning and either Student-Student or Teacher-Student ISC during the review.

## Discussion

Teaching and learning are key processes by which humans build cultural knowledge, enabling the effective transmission of novel and sometimes highly abstracted information (Hasson et al., 2012; Hasson and Frith, 2016). Here, we report the first fMRI study to identify high-level regions of the brain, including regions of the DMN, that are coupled between teachers and students during naturalistic teaching and learning of novel information, and that ultimately predict learning outcomes. Previous work using EEG and fNIRS have the benefit of portability and better temporal resolution, but our use of fMRI allows for full-brain coverage with high spatial resolution. EEG in particular does not allow for measurements from regions of interest in the DMN, including posterior medial cortex and medial prefrontal cortex. Moreover, we employ an extended lecture (32 min) on a single scientific topic, more closely mimicking the typical college-level lecture than the short lessons on unrelated topics that have previously been used. Finally, we recorded the teacher’s brain as she delivered the lessons over multiple sessions, increasing the measurement reliability of the teacher’s neural responses and allowing us to relate both teacher-student and student-student neural coupling to a measure of learning that takes into account both improvement in score and total score.

We observe teacher-student coupling in widespread regions of the brain, extending from visual and auditory areas to high-level areas including DMN. While teacher-student alignment in early sensory regions is likely driven by shared stimulus features, our results suggest that alignment in higher-order regions is driven by shared understanding of lesson content. First, when learning was disrupted by temporally scrambling the lesson, teacher-student coupling was only observed in early visual and linguistic regions, consistent with shared processing of stimulus features (visual input, individual phrases or sentences) on short timescales of a few seconds (Hasson et al., 2008; Lerner et al., 2011; Chen et al., 2015). Teacher-student alignment in DMN and bilateral SPL only emerged during intact learning, consistent with these regions supporting integration of information over longer timescales of minutes (Hasson et al., 2008; Lerner et al., 2011; Honey et al., 2012a; Stephens et al., 2013), as well as high-level context dependent understanding (Honey et al., 2012b; Regev et al., 2013 p.2; Ames et al., 2014; Yeshurun et al., 2017; Nguyen et al., 2019). Second, teacher-student coupling in PMC during the intact lesson significantly predicted learning outcomes, suggesting that students who learn the most are the ones who are most aligned with the teacher. Notably, PMC was also among the regions that only showed teacher-student alignment during intact learning. Our findings are consistent with several recent teacher-student hyperscanning studies: in EEG, teacher-student synchrony in the alpha band (8-12 Hz) is correlated to delayed learning outcomes (Davidesco et al., 2019), while fNIRS have found an association in inferior frontal regions and superior temporal regions (Pan et al., 2018, 2019; Zheng et al., 2018).

Teacher-student coupling was observed in many of the same regions that show coupled responses between speaker and listener during narrative storytelling (Stephens et al., 2010; Dikker et al., 2014; Silbert et al., 2014). These regions include linguistic regions STS and STG extending to the temporal poles, and DMN regions including PMC, mPFC, and dlPFC. However, unlike during narrative storytelling, and in accordance with the scientific nature of our lecture, we also observed teacher-student alignment in bilateral SPL, which has been implicated in mental rotation, geometry, and mathematical symbols (Gogos et al., 2010; Prescott et al., 2010; Zhang et al., 2012; Harvey et al., 2013), as well as the manipulation and rearrangement of information in working memory (Koenigs et al., 2009). We also do not observe teacher-student coupling in bilateral TPJ, a region of the brain that is strongly implicated in mentalizing and theory of mind (Saxe and Kanwisher, 2003; Schilbach et al., 2008; Mars et al., 2012), suggesting that aspects of speaker-listener coupling are content-specific.

Unlike previous studies (Stephens et al., 2010; Silbert et al., 2014; Zadbood et al., 2017; Davidesco et al., 2019), we did not observe a reliable temporal lag between the teacher’s and the listener’s neural responses. The time-locked visual stimulus viewed by both the teacher and the students may have exerted a strong bottom-up, visual signal that was absent in narrative storytelling studies that used auditory-only stimuli. In addition, the low temporal resolution of fMRI and the temporal interpolation procedure for averaging teacher sessions may obscure a temporal lag. Indeed, a recent EEG study with both visual and auditory processing identified a short, 200-ms lag between the teacher and student neural responses (Davidesco et al., 2019). We also did not observe significant differences in the level of neural coupling between the first Lesson video and the second Review video. Based on previous research suggesting that context increases neural coupling, we had hypothesized that the initial lesson would serve to create a shared knowledge base, which would then be reflected in greater student-student and teacher-student alignment during the review than during the lesson. The lack of significant differences between two scans could be due to countervailing differences between the two stimuli. For example, this result may reflect the nature of teaching: a teacher gradually builds common ground with students by connecting novel ideas to basic concepts. Using this “scaffolding” approach, the teacher should effectively guide the student’s learning process to minimize gaps in understanding, thus reducing the difference in coupling between learning and review.

Overall, these findings suggest that neural coupling between teachers and students can be used as an index of learning. While the precise biological processes that give rise to neural alignment has not yet been elucidated, we suggest that neural alignment reflects shared representation of semantic knowledge (Vodrahalli et al., 2018; Cetron et al., 2019; Nastase et al., 2019b). Teaching is therefore a process of building or creating this shared semantic knowledge and aligning representations between students (novices) and teachers (experts). For some concepts (e.g. “net magnetization” or “action potential”), students may be starting from scratch, requiring teachers to build up new concepts from their existing semantic or conceptual knowledge. In other cases, however, teaching may require shifting existing concepts to new contexts. For example, the word “BOLD” in everyday life describes a fearless or daring person or idea. In fMRI, however, “BOLD” refers to the blood-oxygen-level-dependent response. Teachers (or experts) may be able to flexibly shift between conceptual representations of the word “BOLD” while students (or novices) will need to more laboriously shift representations from “fearless” to “fMRI.” Alignment of conceptual knowledge and shared understanding of terms is then reflected in shared neural responses between teachers and students. Joint attention also likely has a role in synchronizing neural responses among individuals (Golumbic et al., 2013; Dikker et al., 2014; Cohen et al., 2018; Regev et al., 2018).

## Limitations and future directions

The role of attention in communication and eliciting shared neural responses in an area of active research (Schmälzle et al., 2015; Cohen and Parra, 2016; Cohen et al., 2018; Regev et al., 2018). Indeed, some have suggested that joint attention to stimulus features during processing of naturalistic stimuli may underlie coupled neural responses (Cohen et al., 2018; Dikker et al., 2019). Decreased Teacher-Student Unscrambled coupling in high-level regions, including DMN, may thus reflect lower attention rather than disrupted learning. However, we also observe that Teacher-Student coupling in sub-regions of the DMN is correlated with learning outcomes. Thus while shared attention is necessary for shared understanding, both like play a role in inducing neural coupling.

Future studies are needed to further explore the building and shifting of shared representations of concepts over time. While we do not observe differences in teacher-student alignment between lesson and review, studies over longer time periods (e.g. over a semester of classes) or using higher temporal resolution imaging coupled with more frequent measures of learning may be able to detect changes in neural alignment reflecting changes in conceptual alignment. Such work may dissect the content of learning by taking advantage of semantic embeddings (e.g., Mikolov et al., 2013). Finally, future studies are needed to further explore the effects of joint attention during interactive learning and teaching using hyperscanning. While the present work explores unidirectional communication and learning, which is increasingly common in the era of online, remote learning, much of learning occurs interactively, in person. During in-person teaching, teachers may be able to adjust their style and explanation following real-time student feedback, enabling better teacher-student alignment and hence learning in contrast to no-feedback conditions.

## Conclusions

In conclusion, this work provides evidence that shared understanding of technical, non-narrative information during teaching and learning is reflected in the coupled neural responses between teachers and students in the DMN and other high-level regions. These findings speak to the flexibility and importance of the DMN in integrating novel information over long time periods and provides insight into how our brains build a shared understanding via communication.

## Acknowledgements

This work was supported by NIH Grants R01MH112566-01 (MN, EM, SN) and 5DP1HD091948-92 (MM, UH) and Princeton ReMatch+ Summer Funding (AC). We thank the entire Hasson Lab for helpful feedback on the analysis and paper.

